# Global Profiling of c-Jun and JunB transcription factor binding sites in an ALK+ ALCL cell line

**DOI:** 10.1101/2022.06.10.495605

**Authors:** Zuoqiao Wu, Mary Nicoll, Farynna Loubich Facundo, Jingxi Zhang, Robert J. Ingham

## Abstract

Anaplastic lymphoma kinase-positive, anaplastic large cell lymphoma (ALK+ ALCL) is a T cell lymphoma which features translocations or inversion involving the *ALK* tyrosine kinase gene, and results in oncogenic fusion proteins (e.g. NPM-ALK). The elevated expression and/or activation of activator protein-1 (AP-1) transcription factors, c-Jun and JunB, is another molecular feature of ALK+ ALCL. c-Jun/JunB transcriptional targets are important in the pathobiology of this lymphoma, and several are also therapeutic targets. To better understand c-Jun/JunB function in ALK+ ALCL, we performed chromatin immunoprecipitation–sequencing experiments in the Karpas 299 ALK+ ALCL cell line to comprehensively identify sites bound by these transcription factors. We identified 13,083 c-Jun and 40,369 JunB binding sites, and ∼60% of sites bound by c-Jun were shared with JunB. Many sites were associated with genes known or predicted to be important in the pathogenesis of ALK+ ALCL. Pathway enrichment analysis of genes associated with both c-Jun and JunB binding sites revealed a significant over-representation for pathways associated with cancer and cell signalling. Furthermore, we identified several c-Jun and JunB binding sites associated with the *NIBAN2*/*FAM129B* gene. FAM129B is a PH domain-containing phosphoprotein that promotes proliferation in multiple cell types. However, while we found that FAM129B knock-down resulted in modest cell cycle alteration in most ALK+ ALCL cell lines, this did not appear to result in a significant proliferation defect. Finally, we found that inhibition of NPM-ALK and MEK/Erk signalling altered FAM129B electrophoretic mobility and decreased phosphorylation of FAM129B on serine residues known to be Erk phosphosites. In summary, this study is the first to globally profile sites bound by c-Jun/JunB in ALK+ ALCL. It reveals novel putative targets for these transcription factors in ALK+ ALCL, and identifies FAM129B as a novel phosphoprotein downstream of NPM-ALK signalling.

## Introduction

Anaplastic lymphoma kinase-positive, anaplastic large cell lymphoma (ALK+ ALCL) is a mature T cell lymphoma that usually presents in children and young adults (1, 2). This lymphoma is named for the presence of chromosomal translocations and inversions involving the *ALK* tyrosine kinase gene, and the t(2;5) translocation with the gene encoding for the nuclear protein, nucleophosmin (NPM), is most common (1-3). These genomic alterations generate oncogenic fusion proteins (e.g. NPM-ALK) with constitutive tyrosine kinase activity, and these fusion proteins activate the Jak/STAT, PI3K, MEK/ERK, and other signalling pathways (3, 4). As well, the elevated expression and/or activation of the Activator Protein-1 (AP-1) family transcription factors, c-Jun and JunB, are characteristic of ALK+ ALCL (5-7). AP-1 family transcription factors consist of members of the Jun (c-Jun, JunB, JunD), Fos (c-Fos, FosB, Fra1, Fra2), ATF, and Maf subfamilies (8-10). These proteins form homo or heterodimers and bind TPA responsive elements (TRE), cAMP response elements (CRE), and variants of these sequences (8-10).

We and others have been interested in elucidating the function of c-Jun and JunB in ALK+ ALCL. siRNA/shRNA knock-down studies have implicated JunB in promoting proliferation in ALK+ ALCL cell lines (5, 11, 12). Studies examining the role of c-Jun in regulating ALK+ ALCL proliferation have come to different conclusions. One study reported that c-Jun knock-down impaired proliferation (13), whereas two others observed no effect on proliferation when c-Jun was knocked-down in ALK+ ALCL cell lines (5, 12). Neither c-Jun nor JunB knock-down in ALK+ ALCL cell lines was found to increase apoptosis (12).

Several c-Jun/JunB transcriptional targets have been identified in ALK+ ALCL, and these are associated with the phenotype and/or linked to pathogenesis in this lymphoma (14). We demonstrated that the serine protease, *Granzyme B* (*GZMB*), a well-known feature of ALK+ ALCL (15, 16), is a JunB transcriptional target in this lymphoma (17), and Granzyme B expression might be a contributing factor for the sensitivity of ALK+ ALCL to chemotherapeutic drugs (18). The gene encoding for the tumour necrosis factor receptor superfamily member, CD30, is also a target of JunB in ALK+ ALCL, and signalling through CD30 also promotes *JunB* transcription (6). Likewise, *Platelet-Derived Growth Factor Receptor-β* (*PDGFR-β*) expression was demonstrated to be promoted by c-Jun and JunB expression in a mouse model of ALK+ ALCL (19). Other transcriptional targets of c-Jun and/or JunB in this lymphoma include genes encoding for the serine/threonine kinases, *Akt1* and *Akt2* (20), and the co-chaperone protein, *Cyclophilin 40* (21). As well, JunB was reported to repress the expression of the DNA helicase, *DDX11*, and in doing so, contributes to chromosome instability in ALK+ ALCL tumour cells (22).

While many functions for c-Jun and JunB in this lymphoma have been defined, the fact that AP-1 binding sites are abundant in the genome (23), suggests that we still only have a cursory understanding of c-Jun/JunB activity in ALK+ ALCL. In addition, these proteins have been reported to have unique (5, 6, 12, 17) and overlapping functions (11, 19, 20) in this lymphoma, but how unique and how much they overlap has not been comprehensively investigated. To address these gaps in knowledge, we performed chromatin immunoprecipitation-sequencing (ChIP-Seq) experiments to identify sites bound by c-Jun and JunB in the Karpas 299 ALK+ ALCL cell line.

## Results

### Identification and characterization of sites bound by c-Jun and JunB

To identify sites bound by c-Jun and JunB in ALK+ ALCL, ChIP-Seq was performed on exponentially growing Karpas 299 cells using c-Jun or JunB specific antibodies. We identified 13,083 sites bound by c-Jun and 40,369 sites bound by JunB **(Fig 1A)**. Fifty-nine percent (7,669) of sites bound by c-Jun overlapped with JunB sites **(Fig 1B)**. Focusing on the most prominent sites bound by c-Jun and JunB, as measured by peak size, we found that slightly less than half of the top 1,000 (and ties) binding sites were shared by c-Jun and JunB **(Fig 1C)**, but this decreased to 25% when the top 100 (and ties) sites were examined **(Fig 1D)**. We also examined whether prominent binding sites for one transcription factor had an overlapping binding site for the other transcription factor. **Fig 1E** shows that ∼75% of the top 1,000 c-Jun binding sites were also bound by JunB, whereas greater than 90% of the top 1,000 sites bound by JunB had a corresponding c-Jun binding site. This overlap was even greater when the top 100 binding sites were examined **(Fig 1E)**. An examination of the sequences associated with the top 1,000 sites bound by c-Jun or JunB revealed a consensus sequence for both transcription factors that conforms to the well-known AP-1 binding TRE motif **(Fig 1F)**. Taken together, these results demonstrate that sites bound by c-Jun and JunB are prevalent in the genome of Karpas 299 cells, and that c-Jun and JunB share a significant number of binding sites.

**Fig 1.**
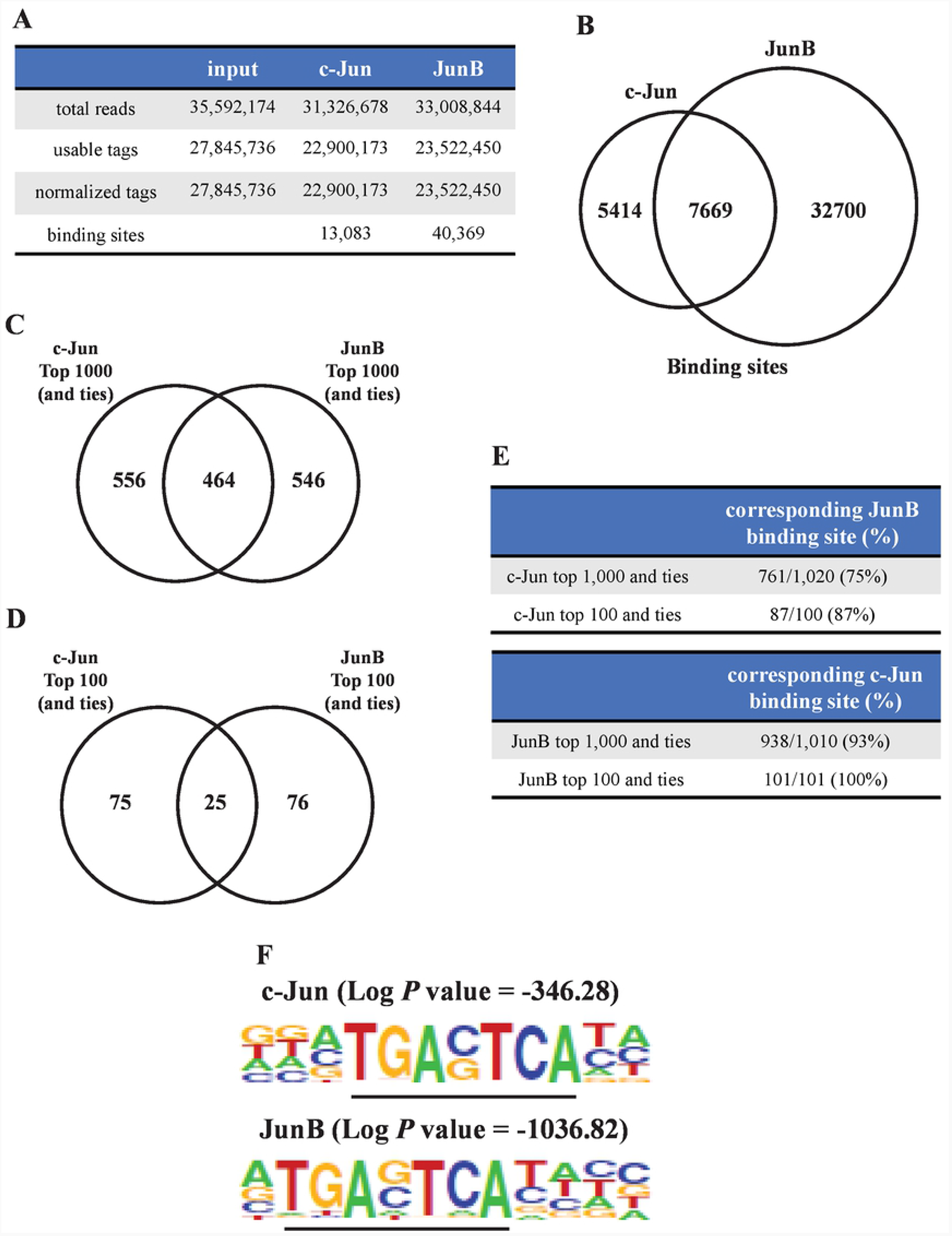
Identification and characterization of sites bound by c-Jun/JunB in Karpas 299 cells. **A**, Summary of the number of reads, tags, and binding sites identified in the ChIP-Seq experiments. **B**, Venn diagram showing the number of binding sites associated with c-Jun alone, JunB alone, or both c-Jun and JunB. Venn diagrams comparing the top 1,000 (and ties) **(C)** or top 100 (and ties) **(D)** c-Jun/JunB binding sites with largest peak size. **E**, Summary of the number (and %) of the top 1,000/100 (and ties) c-Jun or JunB binding sites with a corresponding binding site for the other transcription factor. **F**, Consensus sequence of the top 1,000 c-Jun or JunB binding sites (based on peak intensity) revealed by findMotifsGenome program (v4.8) of the HOMER package (http:homer.ucsd.edu) (24).

### Identification and characterization of genes associated with c-Jun/JunB binding sites

Examining the genomic location of c-Jun/JunB binding sites revealed that the majority of binding sites (84% for c-Jun and 74% for JunB) were located within 10 kb of one or more known genes **(Fig 2A)**. Many binding sites were found within putative promoter regions (−7.5 kb/+2.5 kb from transcriptional start sites) **(Fig 2A)**, and enriched around transcriptional start sites (TSS) **(Fig 2B)**. Moreover, many sites bound by c-Jun/JunB were located near known CpG islands **(Fig 2A)**, which are associated with promoter regions (25). We also investigated where the most prominent sites bound by c-Jun/JunB were located in relation to gene transcriptional start sites. Of the top 1,000 c-Jun binding sites, greater than 30% were within 1 kb of a transcriptional start site of 1 or more genes, and 22% of the top 1,000 JunB binding sites were +/-1 kb from a transcriptional start site **(Fig 2C)**. When the top 100 binding sites were examined, we found that 62% of c-Jun binding sites were within 1 kb of a transcriptional start site **(Fig 2D)**. In contrast, only 17% of the top 100 JunB binding sites were found +/-1 kb from a transcriptional start site **(Fig 2D)**. More of the prominent sites bound by JunB were either not associated with genes, or as especially observed in the top 100 JunB sites, >10 kb away from a transcriptional start site **(Figs 2C** and **D)**.

**Fig 2.**
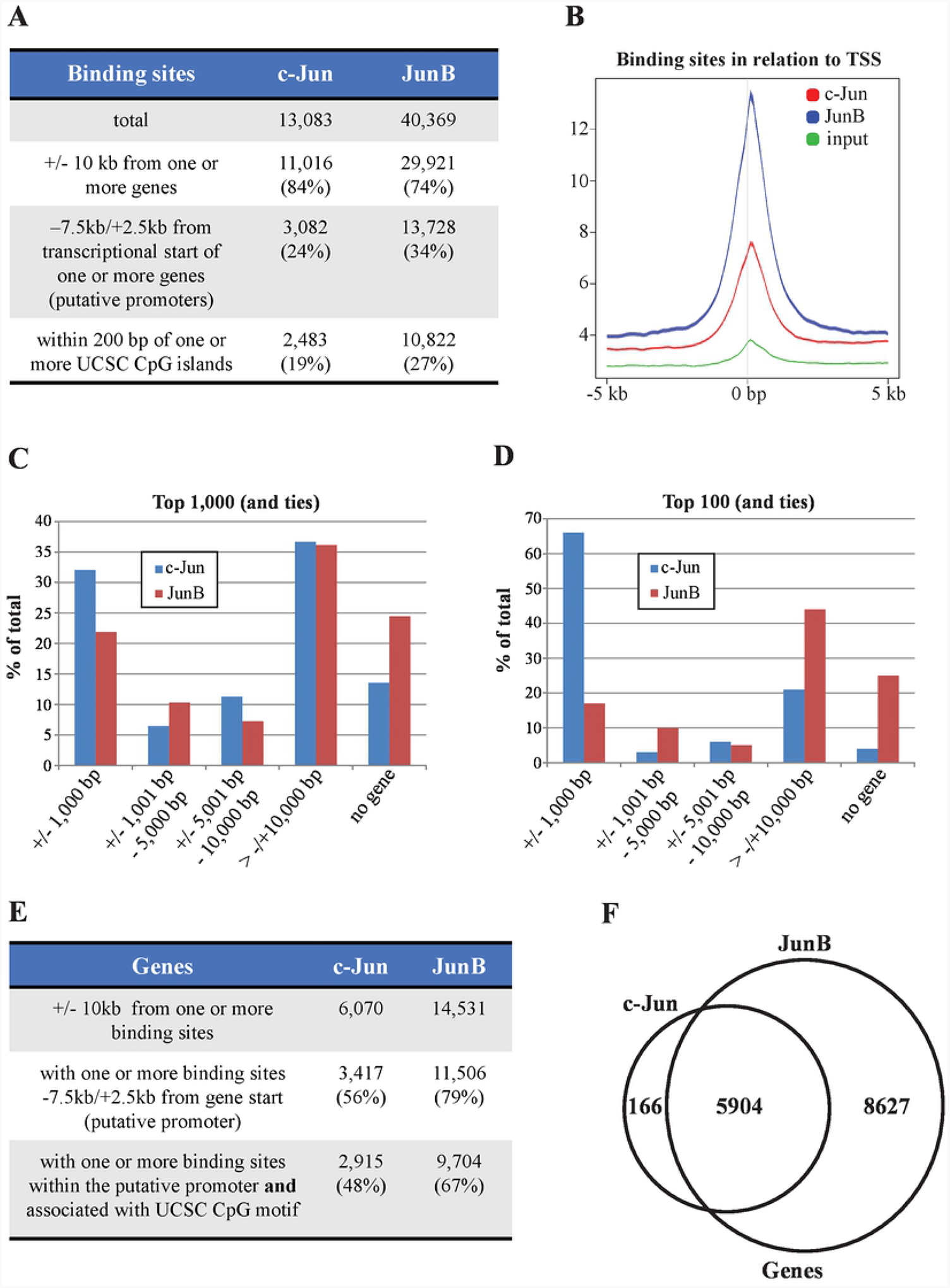
Relationship of c-Jun/JunB binding sites to genes and putative promoter regions. **A**, Summary of the number of c-Jun/JunB binding sites in Karpas 299 that are located within the indicated proximity to genes, transcriptional start sites, and CpG islands. **B**, average plots demonstrating the distribution of binding sites centered on gene transcriptional start sites (TSS). Percentage of top 1,000 **(C)** or top 100 **(D)** binding sites (and ties) within the indicated distance from the transcriptional start site of at least 1 gene. “No gene” indicates that there is no gene within 10 kB of the binding site. **E**, Summary of the number of genes +/-10 kb from c-Jun and JunB binding sites, and the number (and percentage) within putative promoter regions and located 200 bp from a CpG island. **F**, Venn diagram showing the number of genes within 10 kb of c-Jun, JunB, or both c-Jun and JunB binding sites.

An investigation of genes associated with c-Jun/JunB binding sites revealed that 6,070 genes were located within 10 kb of one or more sites bound by c-Jun, whereas 14,531 genes were similarly associated with sites bound by JunB **(Fig 2E)**. The majority of these genes had at least one binding site within putative promoter regions (−7.5 kb/+2.5 kb from transcription start), and many were also located within CpG islands **(Fig 2E)**. Strikingly, 97% of genes within 10kb of c-Jun binding sites also had a JunB binding site; albeit these were not always the binding site(s) **(Fig 2F)**.

We next examined individual genes associated with c-Jun/JunB binding sites. Sites associated with *GZMB, CD30/TNFRSF8, LGALS1/Galectin-1* (26), *CCNA2/Cyclin A2* (11), and other genes previously shown or suggested to be transcriptional targets of AP-1 proteins in ALK+ ALCL were identified **(Fig 3A; S1 Table)**. Focusing exclusively on genes with binding sites +/-1,000 bp from putative transcriptional start sites revealed several genes prominently expressed and/or important in the pathobiology of ALK+ ALCL, but not previously described to be regulated by AP-1 proteins. These include the transcription factors, *STAT3* (27, 28) and *IRF4* (29), the pro-survival Bcl-2 family member, *MCL1* (30, 31), and the cytokine, *IL21* (32) **(Fig 3B; S2 Table)**. Moreover, some of the most prominent binding sites were associated with genes that have not been investigated in ALK+ ALCL, but could play important roles in this lymphoma **(Fig 3C)**. These genes included the ABC transporter, *ABCA3*, whose protein product is associated with drug resistance in lung cancer (33) and chronic myelogenous leukemia (34, 35), and which mediates exosome release to protect B cell lymphomas from anti-CD20 therapy (36). As well, c-Jun and JunB binding sites were associated with the gene encoding for the calcium binding protein, S100A10, which performs multiple functions in cancer including mediating chemoresistance (37). Taken together, our data show that many c-Jun/JunB binding sites are located within putative promoter regions, and the majority of genes associated with c-Jun binding sites are also associated with JunB binding sites. In addition, binding sites are associated with genes known or predicted to be important in the pathology of ALK+ ALCL.

**Fig 3.**
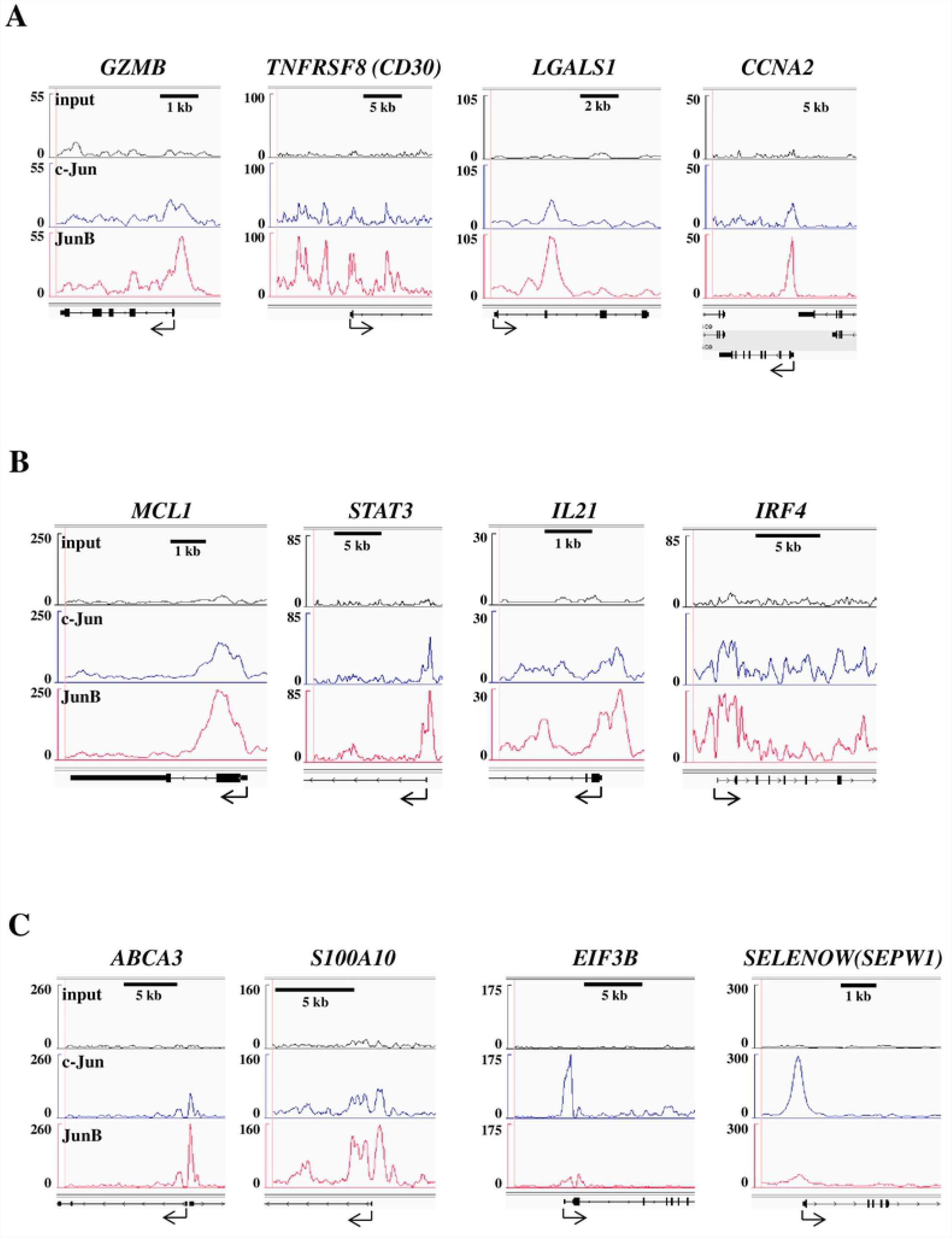
Sites bound by c-Jun/JunB are associated with genes known or predicted to be important in the pathobiology of ALK+ ALCL. Integrated genome viewer (IGV) images showing c-Jun/JunB binding sites associated with: known AP-1 transcriptional targets in ALK+ ALCL (**A**), genes important in the pathogenesis of ALK+ ALCL but not described as AP-1 transcriptional targets (**B**), or the most prominent c-Jun/JunB binding sites (**C**). “input” represents input DNA

### c-Jun/JunB bindings sites are enriched in genes associated with cancer and cellular signalling

We next examined whether specific pathways or cellular activities were enriched for in genes associated with c-Jun/JunB binding sites. Utilizing the g:Profiler on-line tool (38, 39), we searched for enriched pathways in the Gene Ontology (GO) Biological Process (40, 41), Reactome (42), and the Kyoto Encyclopedia of Genes and Genomes (KEGG) (43) pathway databases. Few pathways were significantly enriched for the small number of genes exclusively associated with c-Jun binding sites **(Fig 4A; S3 Table)**. Surprisingly, genes associated with both c-Jun and JunB binding sites were enriched for considerably more pathways than genes associated with JunB alone **(Fig 4A; S3 Table)**. Thi**s** despite the former totalling only ∼70% of the number of genes associated with JunB binding sites alone. A closer examination of KEGG pathways enriched for genes with both c-Jun and JunB binding sites, revealed enriched pathways related to cancer, signalling pathways, and cellular processes whose dysregulation is associated with cancer **(Figs 4B** and **C; S3 Table)**. Thus, genes associated with both c-Jun and JunB binding sites show particular enrichment for pathways that are either known or likely to contribute to the pathobiology of this lymphoma.

**Fig 4.**
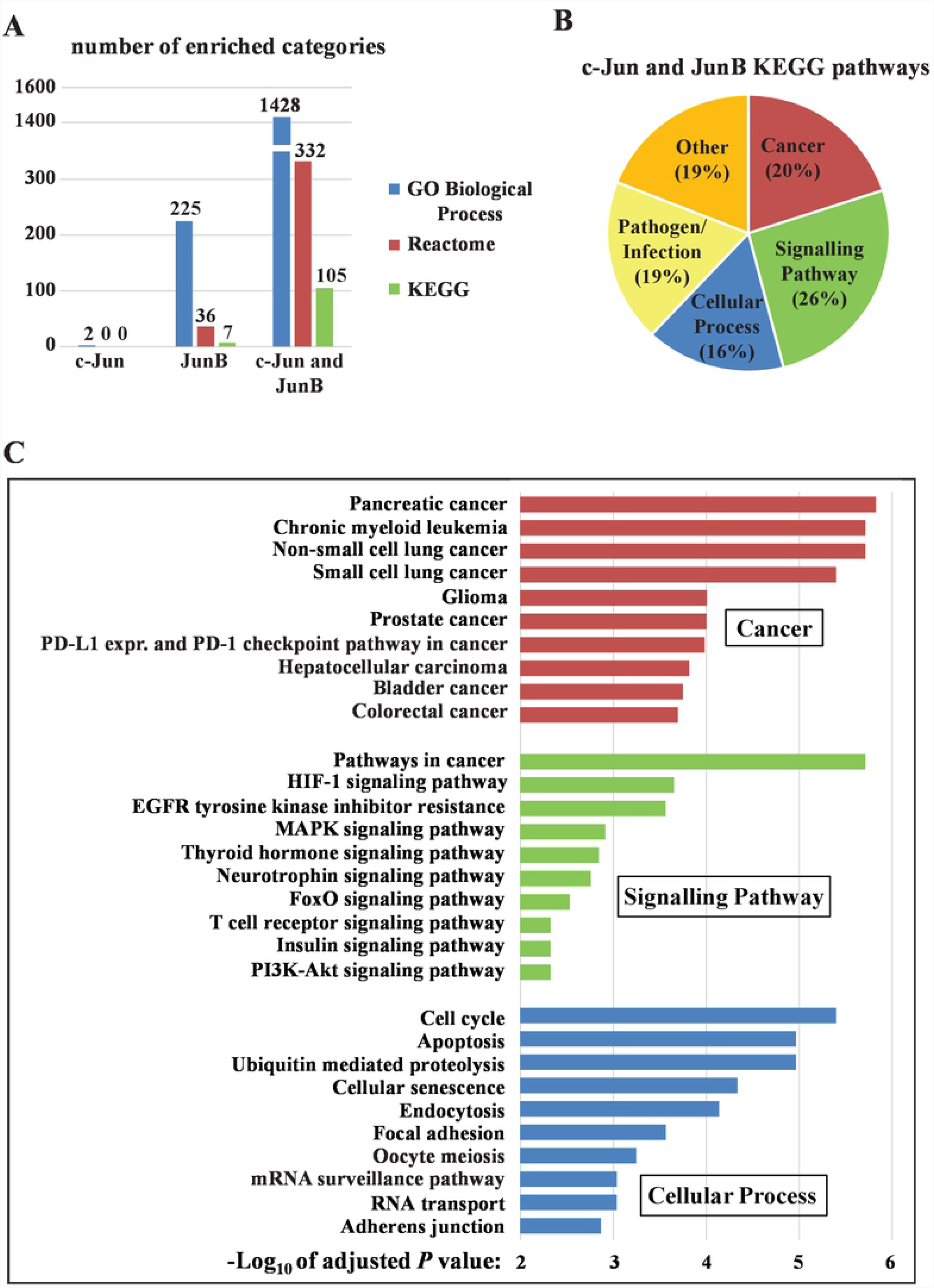
Genes associated with both c-Jun and JunB binding sites are significantly enriched for biological pathways and processes important in cancer. **A**, Comparison of the number of Gene Ontology (GO) Biological Process (blue), Reactome (red), and KEGG (green) pathways associated with genes (+/-10 kb) from c-Jun, JunB, or both c-Jun and JunB binding sites. The numbers on top of each bar indicate the total number of identified enriched pathways with an adjusted *P* ≤ 0.05. **B**, Categories (and percentage of total) of enriched KEGG pathways associated with genes with both c-Jun and JunB binding sites. **C**, The top 10 enriched pathways, based on Log_10_ of adjusted *P* value, from the “Cancer” (red), “Signalling Pathways” (green), and “Cellular Process” (blue) categories in **B**. The complete list of enriched KEGG pathways can be found in **S3 Table**.

### *NIBAN2/FAM129B* is associated with multiple c-Jun/JunB binding sites in Karpas 299 cells

Our ChIP-Seq experiments revealed many genes that were not only potentially novel transcriptional targets of c-Jun and/or JunB in ALK+ ALCL, but could also contribute to the pathogenesis of this lymphoma. Among these genes, we were particularly intrigued with *NIBAN2*, which is alternatively referred to as *MINERVA*, or more commonly, *FAM129B*. FAM129B is a Pleckstrin Homology (PH) domain-containing protein that was first identified as an Erk substrate, and Erk-mediated phosphorylation of FAM129B was shown to be important for invasion in melanoma (44). FAM129B is also substrate for the Epidermal Growth Factor Receptor (EGFR) receptor tyrosine kinase in glioblastoma cell lines, and tyrosine phosphorylated FAM129B binds the Ras GTPase (45). When FAM129B binds Ras, this blocks p120-RasGAP from binding and inactivating Ras. This results in increased Ras/MEK/Erk/β-catenin signalling which promotes proliferation and invasion in glioblastoma cell lines (45). Given that FAM129B is an important phosphoprotein in other cancers, we felt it was an intriguing protein to investigate in ALK+ ALCL.

We identified a number of c-Jun/JunB binding sites associated with *FAM129B* **(Fig 5A)**, and this included a prominent site recognized by both c-Jun and JunB with a consensus TRE site ∼150bp from the transcriptional start site of the larger *FAM129B* isoform **(Fig 5B)**. FAM129B protein expression was evident in all ALK+ ALCL cell lines tested, with higher and lower molecular weight species evident **(Fig 5C)**.

**Fig 5.**
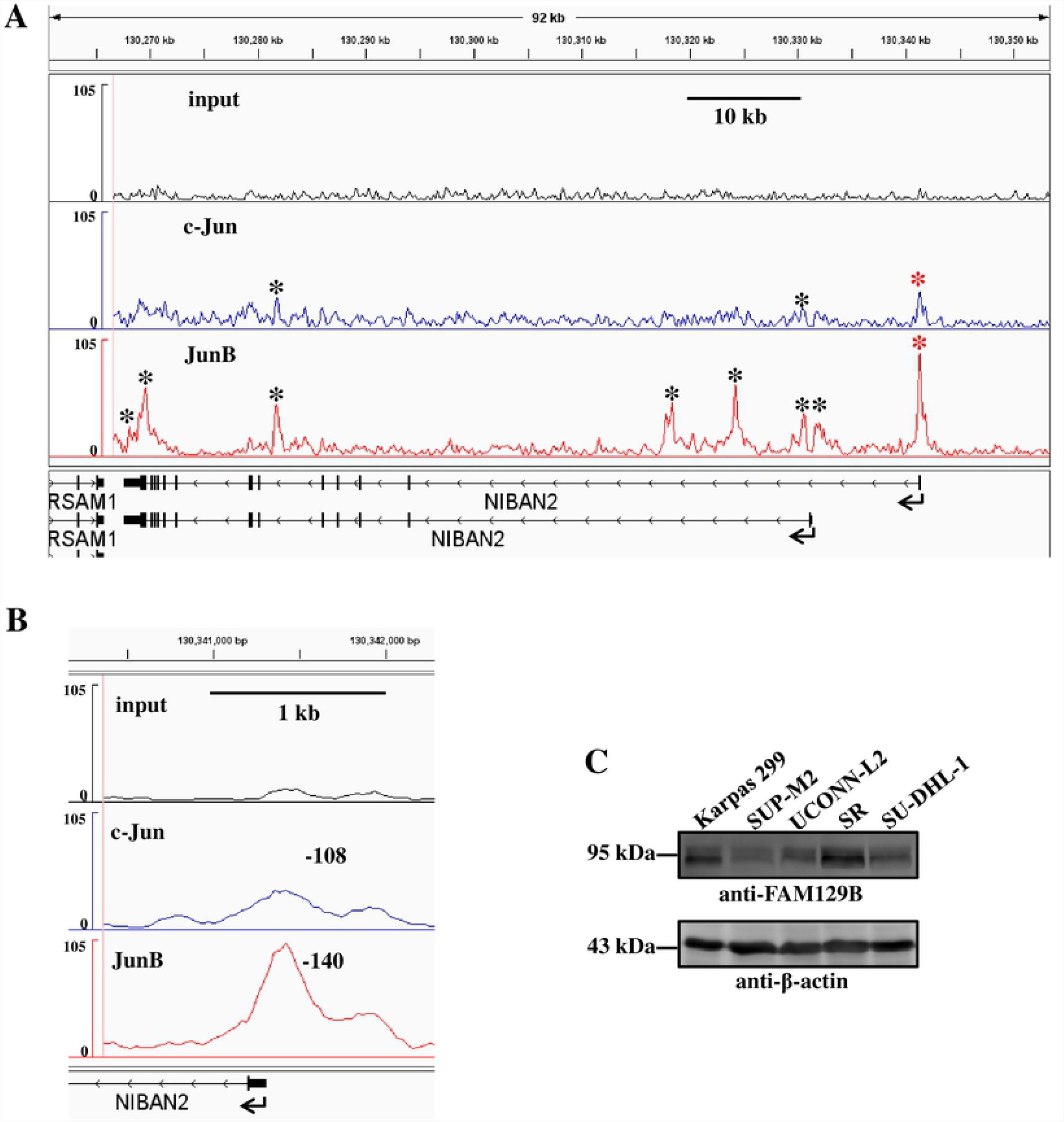
c-Jun and JunB binding sites are associated with *FAM129B*. **A**, Integrated genome viewer (IGV) image showing c-Jun/JunB binding sites (denoted by asterisks) in relationship to *FAM129B* transcripts. The overlapping c-Jun/JunB binding sites denoted by red asterisks were the most prominent binding site and contains a consensus AP-1 TRE motif. **B**, Integrated genome viewer (IGV) image illustrating the prominent overlapping c-Jun/JunB binding site 5’ of the transcriptional start site of the larger *FAM129B* transcript (indicated by a red asterisk in **A**). **C**, Western blotting experiments demonstrating FAM129B protein levels in ALK+ ALCL cell lines. Molecular mass markers are indicated to the left of blots.

We next examined whether the identified TRE site within the prominent c-Jun/JunB binding site ∼150bp from the transcriptional start site of the larger *FAM129B* isoform **(Fig 5B)** was able to bind AP-1 proteins and promote *FAM129B* transcription. To address the former, we performed an electrophoretic mobility shift assay (EMSA) to examine whether JunB could bind a 21mer biotinylated probe based on the sequence surrounding the putative TRE site. A protein, or proteins, within Karpas 299 cells nuclear extracts was able to bind this probe, and binding was inhibited with a 50 molar excess of unlabelled probe, but not an excess of unlabelled probe with a mutation in the TRE motif **(Fig 6A)**. Similarly, binding was observed with nuclear extracts from additional ALK+ ALCL cell lines, and probe/protein complexes were super-shifted by the addition of an anti-JunB Ab demonstrating that JunB was a component of these probe/protein complexes **(Fig 6B)**. To evaluate whether the TRE site was important for *FAM129B* transcription, we generated a luciferase promoter construct consisting of nucleotides +50 to -755bp (relative to the transcriptional start site) of the larger *FAM129B* isoform **(Fig 5A)**. We also generated a similar construct with a mutation in the TRE site. Compared to the *FAM129B* construct, the *FAM129B* TRE mutant construct exhibited reduced activity in Karpas 299 cells **(Fig 6C)**, and ectopic expression of JunB in the DG75 Burkitt lymphoma cell line enhanced luciferase activity from the *FAM129B* construct **(Fig 6D)**. However, the *FAM129B* construct exhibited less activity than our previously described *GZMB* luciferase AP-1 reporter construct (17), and mutation of the TRE motif in the *FAM129B* construct had a more modest impact on luciferase activity than the same mutation in the *GZMB* reporter construct **(Fig 6C)**. Since JunB binding sites associated with *FAM129B* were more prevalent and prominent than those for c-Jun **(Figs 5A** and **B)**, we examined whether knock-down of JunB was sufficient to reduce FAM129B expression. Knock-down of JunB in Karpas 299 cells with two distinct JunB shRNAs resulted in no significant difference in FAM129B protein levels **(Fig 6E)**. In contrast, we observed decreased FAM129B expression when JunB was knocked-down in SUP-M2 cells, and FAM129B levels correlated with the degree of JunB knock-down. These results confirm that JunB binds to the TRE site located upstream of the larger FAM129B isoform; however, this site, and surrounding sequence, were only modestly able to promote luciferase reporter activity in Karpas 299 cells. Nonetheless, at least in SUP-M2 cells, the silencing of JunB was sufficient to reduce FAM129B expression.

**Fig 6.**
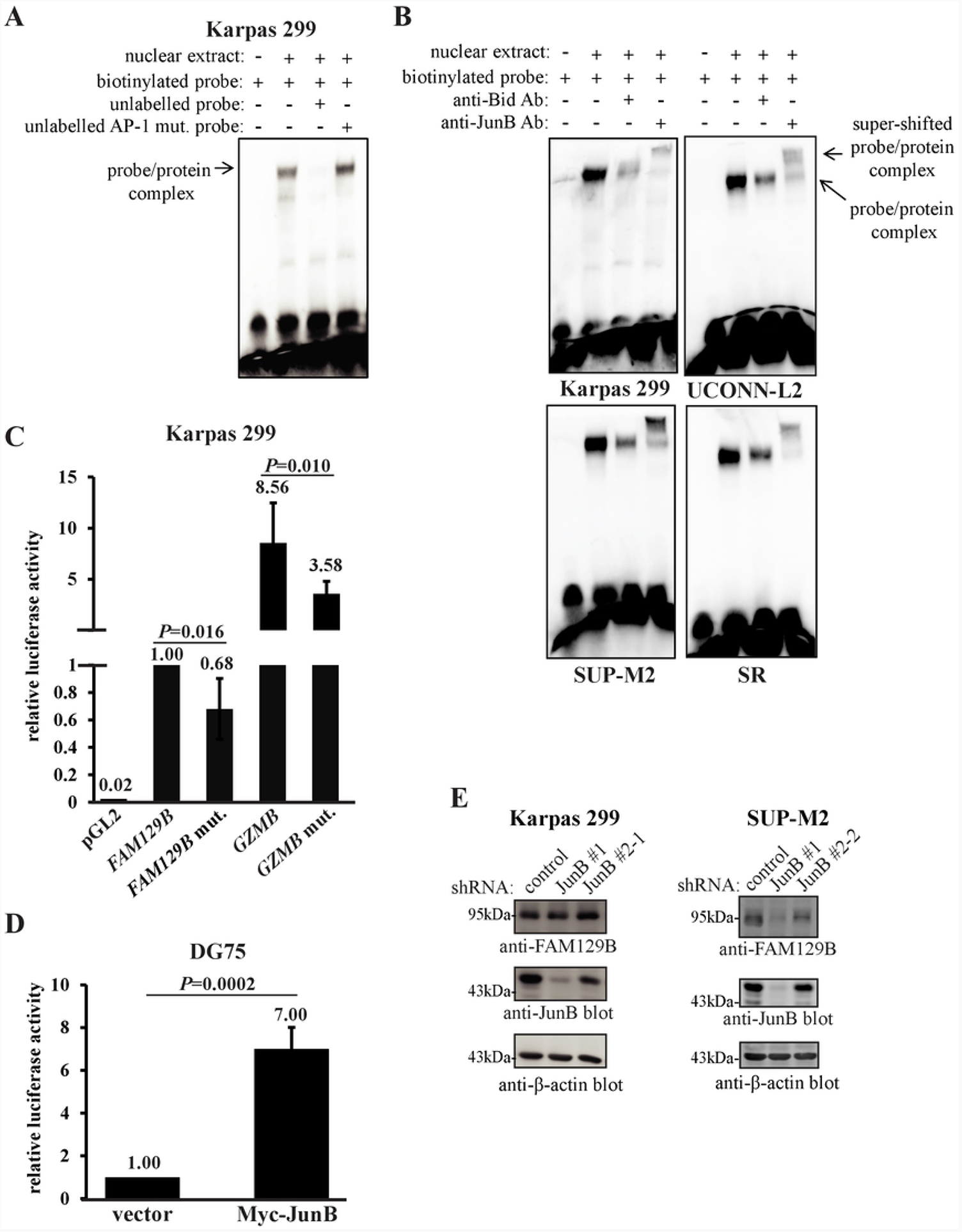
JunB binds the TRE motif within *FAM129B* promoter and modestly promotes *FAM129B* transcriptional activity. **A** and **B**, EMSAs showing the appearance of probe/protein complexes when nuclear extracts from the indicated cell lines were incubated with a 21 mer biotinylated probe based on the TRE site found within the c-Jun/JunB binding site in the *FAM129B* promoter illustrated in **Fig 5B**. In **A**, a 50 molar excess of unlabelled probe (lane 3) or unlabelled probe with a mutation in the TRE site (lane 4) was included in reactions to determine the importance of the TRE site for formation of the probe/protein complex. In **B**, the addition of the anti-JunB or anti-Bid (isotype control) to reactions was included to demonstrate the presence of JunB in the probe/protein complexes. **C**, Karpas 299 cells were transfected with the indicated firefly luciferase reporter constructs (in pGL2 basic) and a constitutive *Renilla* luciferase plasmid. Firefly and *Renilla* luciferase activity was measured 24h post-transfection. The ratio of Firefly to Renilla activity was determined and the results were expressed relative to the pGL2 *FAM129B* (*FAM129B*) which was arbitrarily set at 1. The results shown represent the average and standard deviation of 5 independent experiments. Statistics represent paired, one-tailed *t* tests comparing activity of the wt and corresponding mutant constructs. The empty vector, pGL2 basic (pGL2), was included as a negative control. **D**, DG75 B lymphoma cells were transfected and treated as in **C** with the addition that pcDNA 3.1A Myc-JunB (Myc-JunB) or vector alone (vector) were included in the transfection mixture. The results were expressed relative to the vector-transfected cells which were arbitrarily set at 1. The results show the average and standard deviation of 3 independent experiments. Statistics represent a paired, one-tailed *t* tests. **E**, Western blots demonstrating the expression of FAM129B in Karpas 299 (left) or SUP-M2 (right) cells expressing either control or JunB shRNAs. The anti-β-actin blots are reprobes and illustrate protein levels in the indicated lysates. Molecular mass markers are indicated to the left of blots.

### FAM129B knock-down in most ALK+ ALCL cell lines was associated with a modest alteration in the cell cycle

FAM129B has been implicated in promoting proliferation in glioblastoma (45) and non-small cell lung carcinoma cell lines (46). Therefore, we reduced FAM129B expression in four ALK+ ALCL cell lines using two distinct FAM129B shRNAs **(Fig 7A)**, and performed BrdU/7-AAD double staining to analyze the cell cycle distribution of control and FAM129B shRNA-expressing cells **(S1 Fig)**. We found that FAM129B knock-down, with both shRNAs, moderately decreased the percentage of cells in S phase and increased the percentage in G_0_/G_1_ phase in most cell lines **(Fig 7B)**. However, growth curve experiments performed on Karpas 299 and SUP-M2 cells either did not or inconsistently showed that FAM129B knock-down resulted in a growth defect **(results not shown)**. Therefore, while FAM129B knock-down in most ALK+ ALCL cell lines resulted in a statistically significant alteration in the cell cycle, this alteration did not appear to substantially affect the growth rate.

**Fig 7.**
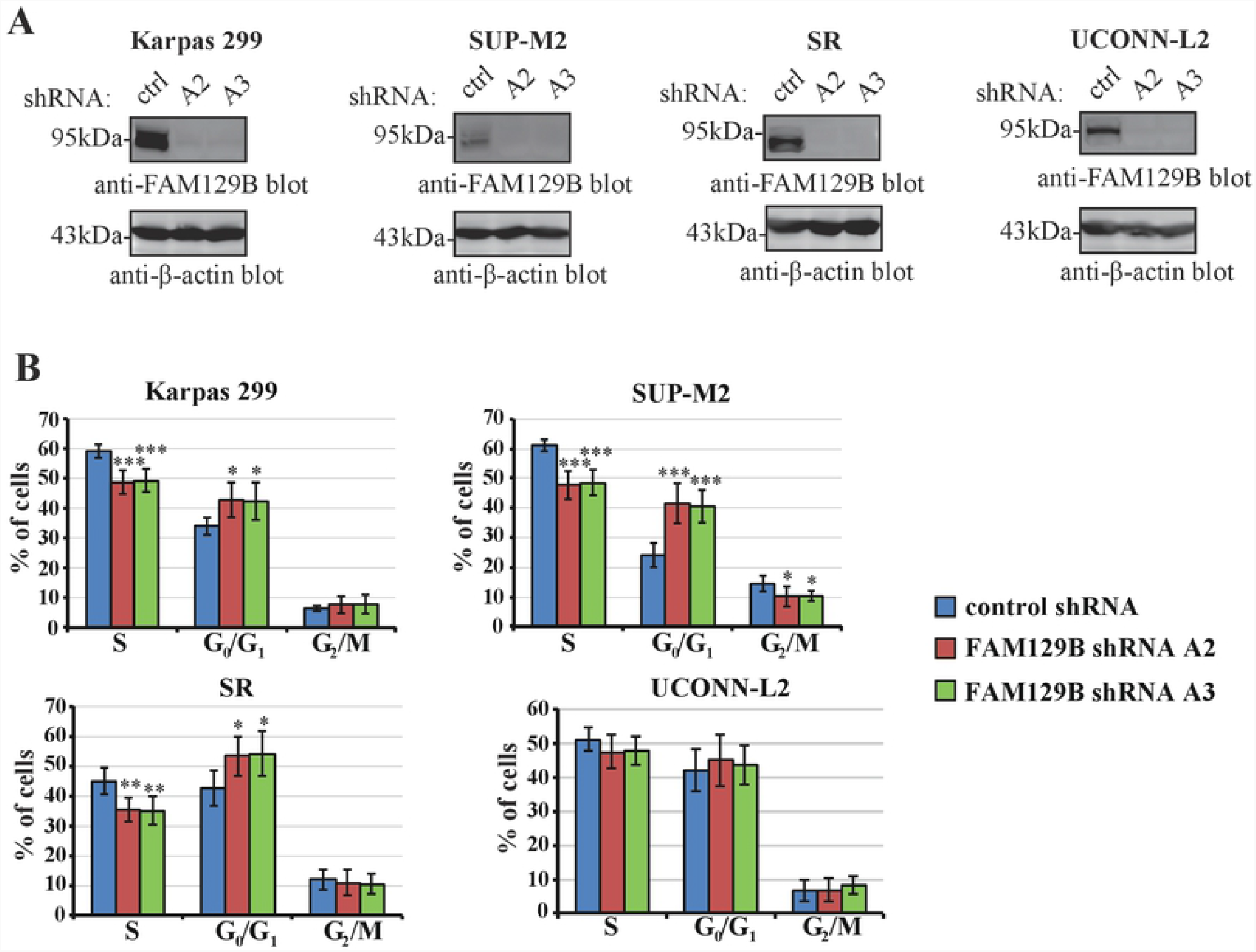
FAM129B knock-down in most ALK+ ALCL cell lines was associated with a modest cell cycle alteration. **A**, Western blots demonstrating the expression of FAM129B in the indicated cell lines expressing either control or the FAM129B shRNAs. The anti-β-actin blots demonstrate protein levels in the indicated lysates. Molecular mass markers are indicated to the left of blots. **B**, Summary of cell cycle distribution from BrdU/7-AAD labelling data in the indicated cell lines expressing either control or FAM129B shRNAs. The results represent the average and standard deviation of seven (Karpas 299), six (SUP-M2), seven (SR), or four (UCONN-L2) independent experiments from at least two independent infections. In all cases, *P* values were obtained by performing ANOVA with a Tukey *post-hoc* test. *, *P* ≤ 0.05; **, *P* ≤ 0.01; ***, *P* ≤ 0.001.

### FAM129B is phosphorylated by Erk in an NPM-ALK-dependent manner in ALK+ ALCL

The fact that we observed multiple molecular weight FAM129B species led us to investigate whether FAM129B might be post-translationally modified in ALK+ ALCL. FAM129B is tyrosine-phosphorylated by the epidermal growth factor receptor (EGFR) (45), and phosphorylated on multiple serine residues in response to Raf/MEK/Erk signalling in melanoma cell lines (44). We observed reduced FAM129B electrophoretic mobility in Karpas 299 cells treated with the ALK tyrosine kinase inhibitor, Crizotinib **(Fig 8A)**. Similar results were observed in SUP-M2 cells **(Fig 8B)**. While we found no evidence that FAM129B was tyrosine phosphorylated in Karpas 299 cell lines (**results not shown**), ALK inhibition resulted in decreased phosphorylation of FAM129B on serines 679 and 683 which are regulated by MEK/Erk signalling in melanoma cell lines (44) **(Fig 8C)**. To confirm that FAM129B electrophoretic mobility was modulated by MEK/Erk signalling in ALK+ ALCL, treatment of Karpas 299 and SUP-M2 cells with the MEK inhibitor, U0216, resulted in increased FAM129B electrophoretic mobility and decreased phosphorylation of FAM129B on serines 679 and 683 **(Fig 8D** and **E)**. To investigate whether NPM-ALK signalling is sufficient to alter FAM129B electrophoretic mobility and serine 679/683 phosphorylation, and to determine whether this is MEK/Erk dependent, we utilized HEK 293P cells expressing a doxycycline-inducible NPM-ALK (47). Induction of NPM-ALK expression in these cells resulted in decreased FAM129B electrophoretic mobility as well as increased phosphorylation of FAM129B on serines 679/683 **(Fig 8F)**. Moreover, treatment with U0126 partially abolished the FAM129B band shift and serine 679/683 phosphorylation in response to NPM-ALK induction **(Fig 8F)**. Taken together, these results demonstrate that NPM-ALK signalling regulates FAM129B serine phosphorylation via MEK/Erk signalling.

**Fig 8.**
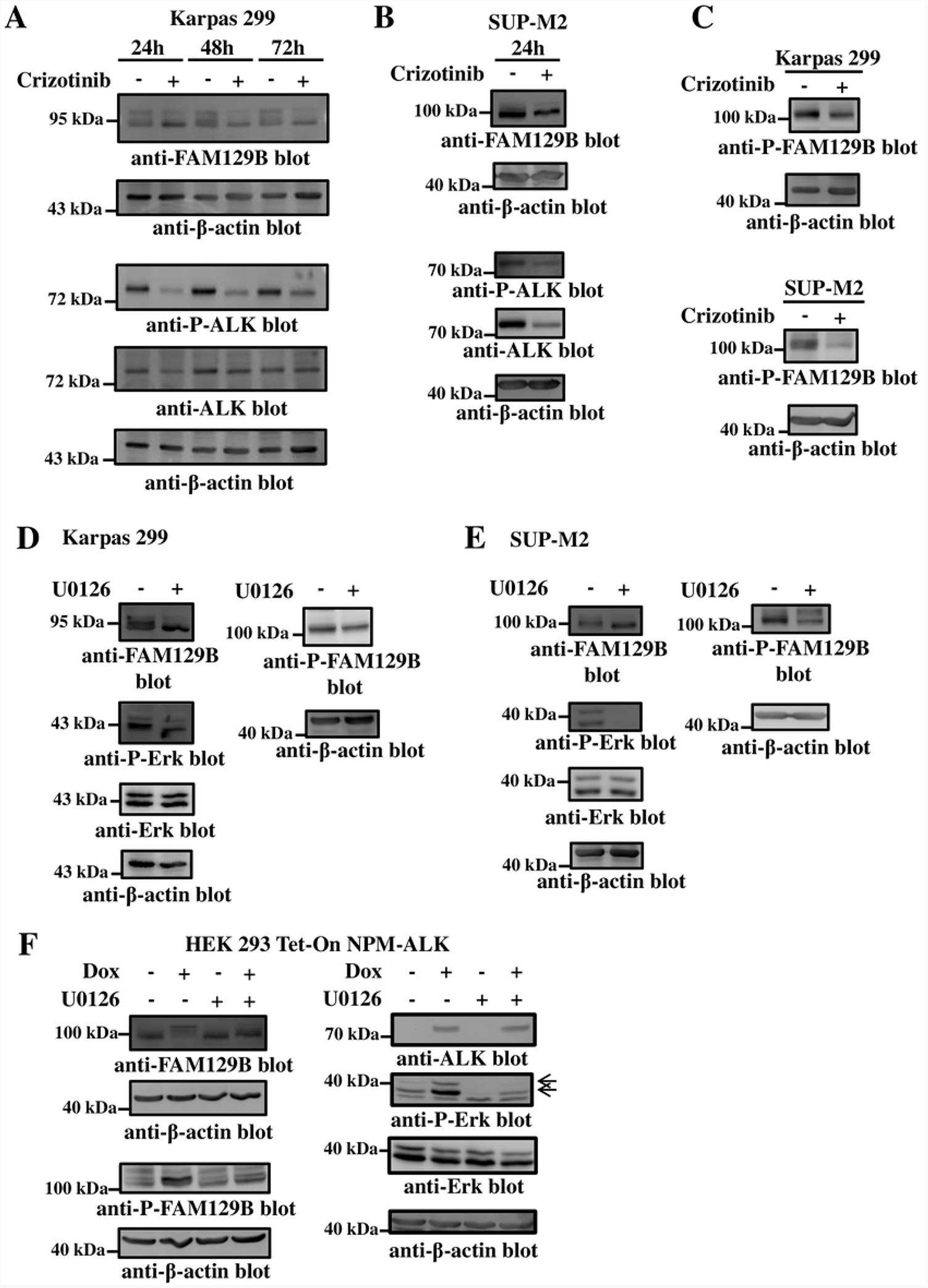
FAM129B is phosphorylated by Erk in an NPM-ALK-dependent manner. **A**, Karpas 299 cells were treated with the 75 nM of Crizotinib for the indicated times. Lysates were then western blotted with the indicated antibodies. **B**, SUP-M2 cells were treated for 24 h with 150 nM Crizotinib prior to lysis, and lysates were probed with the indicated antibodies. **C**, Karpas 299 (above) or SUP-M2 cells (below) were treated with 75 or 150 nM of Crizotinib, respectively for 24 h before being lysed. Lysates were subsequently probed with P-FAM129B or β-actin antibodies. Karpas 299 **(D**) or SUP-M2 **(E)** cells were treated for 6 h with 10 μM U0126 (+) or DMSO alone (−) before being lysed. Lysates where then probed with the indicated antibodies. **F**, HEK 293 Tet-On NPM-ALK cells were either left untreated or treated with 5 µg/mL of Doxycycline (Dox.) for 24 h at which time cells were either treated with 10 μM U0126 or DMSO control. Lysates were then western blotted with the indicated antibodies. Arrows indicate P-Erk. For all blots, anti-β-actin blots illustrate protein levels in the lysates. Molecular mass markers are indicated to the left of blots.

## Discussion

The elevated expression of c-Jun and JunB are well-recognized features of ALK+ ALCL, and these transcription factors regulate the expression of many genes that characterize this lymphoma. In this study, we have utilized ChIP-seq to generate a more complete picture of genomic binding sites occupied by c-Jun and JunB in ALK+ ALCL, and by extension, their putative transcriptional targets. Although our ChIP-Seq was limited to a single cell line, our findings have revealed important new information regarding the activities of c-Jun and JunB in ALK+ ALCL that will be complemented by future ChIP-Seq studies expanded to addition cell lines and/or patient samples.

We identified 40,369 sites bound by JunB, and slightly less than a third as many sites bound by c-Jun in Karpas 299 cells **(Fig 1)**. Moreover, peak intensities for shared binding sites were generally greater for JunB than c-Jun (see **Figs 3** and **5** as examples). While this may partly be due to differences in the quality of antibodies used for the ChIP-Seq experiments and we may be underestimating c-Jun binding sites, other studies have also suggested a more important role for JunB than c-Jun in ALK+ALCL. Firstly, quantitative RT-PCR experiments found that JunB is expressed at higher levels in ALK+ ALCL cell lines than c-Jun (5). Secondly, two knock-down studies identified a more significant role for JunB in the regulation of proliferation (5, 12). Nonetheless, we observed significant overlap between sites bound by c-Jun and JunB **(Fig 1B)**, particularly when the most prominent sites were examined **(Fig 2)**. This suggests that there is considerable intersection between cellular activities and gene targets regulated by c-Jun and JunB, and perhaps other AP-1 family members, and that this redundancy could result in phenotypes not being observed in knock-down studies where only one of the proteins is reduced in expression.

Another intriguing observation from this study was that the location of prominent binding sites differed between c-Jun and JunB. Whereas prominent c-Jun binding sites were more likely to be located within 1,000 bp of a transcriptional start site, especially for the top 100 binding sites, JunB binding sites with highest peak intensity were more likely to be either located greater than 10,000 bp from a gene transcriptional start site or not within 10 kb of a known gene **(Figs 2C** and **D)**. This argues that prominent c-Jun binding sites may preferentially be present at proximal promoters, whereas many prominent JunB sites could be located in putative enhancers. While the majority of studies have characterized the importance of c-Jun/JunB binding sites acting close to transcriptional start sites in ALK+ ALCL ((6, 17, 21) as examples), it is well established that AP-1 proteins also function at enhancers and can regulate gene expression at sites distant from promoters (48).

Our study identified genes previously described to be regulated by AP-1 proteins in ALK+ ALCL such as *CD30* (6), *GZMB* (17), *PDGFRB* (19), and *AKT* (20) **(Fig 3A)**. In addition, sites bound by these transcription factors were associated with other genes known to be important in the pathogenesis of ALK+ ALCL which could represent novel c-Jun/JunB-regulated genes in this lymphoma **(Fig 3B)**. Considering only genes with a binding site +/-1kb from the transcriptional start site, we identified the *STAT3* transcription factor which promotes pro-proliferation and pro-survival signalling in ALK+ ALCL (27, 28), as well as the Bcl-2 family protein, *MCL1*, which is highly expressed in ALK+ ALCL (30, 31) and implicated in promoting survival (49, 50). We also identified binding sites associated with *BATF*, a member of the extended AP-1 family, which was shown to regulate growth, survival, and the expression of genes associated with T_H_17 cells in ALK+ ALCL (51). Finally, we identified binding sites associated with the *IL21* and *IL21R* which activates Jak/STAT signalling to promote cell growth in ALK+ ALCL (32). Thus, while the regulation of these genes by c-Jun and/or JunB still needs to be confirmed, our observations suggest that these transcription factors regulate additional genes with important roles in ALK+ ALCL. Moreover, since we focused our analysis on genes with binding sites +/-1 kb from a transcriptional start, including genes with binding sites outside of this distance further expands the potential roles for these transcription factors in this lymphoma.

The full scope of the pathways and processes putatively regulated by c-Jun and JunB in ALK+ ALCL was revealed in our pathway enrichment analysis; in particular, we were intrigued by the number of enriched pathways for genes associated with both c-Jun and JunB binding sites **(Fig 4 and S3 Table)**. This argues that many roles that c-Jun and JunB perform in this lymphoma are shared, and these shared roles include pathways that would be predicted to benefit this lymphoma **(Figs 4B** and **C)**. However, it remains to be determined whether these genes are truly regulated by c-Jun/JunB, and if so, whether these transcription factors equally contribute to the transcription of these genes.

In this study, we characterized one gene associated with prominent c-Jun and JunB binding sites, *FAM129B* **(Figs 5A** and **B)**. Intriguingly, *FAM129B* was identified in a microarray study as a gene highly expressed in ALCL patients (both ALK+ and ALK-) compared to patients with Peripheral T cell lymphoma-not otherwise specified (PTCL-NOS) and Angioimmunoblastic T cell lymphoma (AITL) (52). However, *FAM129B* was not amongst the most prominent up-regulated genes when ALK-ALCL alone was compared to PTCL-NOS and AITL. We determined that the consensus TRE motif within the prominent c-Jun/JunB binding site immediately upstream of the transcriptional start site of the larger *FAM129B* isoform was able to bind JunB **(Figs 6A** and **B)**. Surprisingly, we found that a luciferase reporter construct including this site displayed modest activity in Karpas 299 cells, but this activity was decreased when the TRE site was mutated **(Figs 6C** and **D)**. This contrasted with our previously described *GZMB* reporter construct (17) which exhibited significant luciferase promoter activity in Karpas 299 cells **(Fig 6C)**. It could be that this site functions as an enhancer that promotes transcription of the more distal, smaller *FAM129B* isoform **(Fig 5A)**, but this will require further examination. Nonetheless, our results show that, at least in SUP-M2 cells, JunB knock-down was sufficient to reduce FAM129B expression **(Fig 6D)**. Whether this involves the site we characterized, other c-Jun/JunB binding sites **(Fig 5A)**, or indirect regulation by JunB is unknown.

While we found that FAM129B knock-down resulted in a consistent, statistically significant cell cycle alteration in most ALK+ ALCL cell lines, these changes were modest **(Fig 7)**. The greatest effect was observed in SUP-M2 cells **(Fig 7B)**, but we could not convincingly demonstrate that FAM129B knock-down translated to a significant growth defect even in these cells **(results not shown)**. Thus, given these results, we conclude that FAM129B does not significantly influence proliferation in ALK+ ALCL and may perform another function in this lymphoma. FAM129B promotes invasion in melanoma (44, 53), glioblastoma (45), and non-small lung carcinoma (46) cell lines. As well, FAM129B counters oxidative stress in multiple cancer types through activation of the anti-oxidative transcription factor, Nrf2 (53-55). These are intriguing possibilities for how FAM129B could function in ALK+ ALCL.

While the downstream consequences of FAM129B signalling in ALK+ ALCL remain elusive, we show that FAM129B is serine phosphorylated by MEK/Erk signalling in a NPM-ALK-dependent manner in this lymphoma. NPM-ALK and MEK/Erk kinase activity was required for a higher molecular weight FAM129B species, and phosphorylation of FAM129B on serines 679 and 683 which are known Erk phosphosites (44) **(Fig 8)**. Serine/threonine phosphorylation is critical for FAM129B to mediate invasion in melanoma cell lines, and U0126 treatment was found to re-localize FAM129B from the membrane to the cytosol (44, 53). Cytosolic FAM129B binds Keap1, and thereby prevents Keap1 from binding Nrf2, and targeting Nrf2 for ubiquitin-mediated degradation (53). Thus, MEK/Erk-mediated phosphorylation of FAM129B modulates FAM129B activity, and our observations suggest that this might represent a mechanism to regulate FAM129B activity in ALK+ ALCL.

In summary, this study provides the first comprehensive overview of sites occupied by c-Jun and JunB in the genome of an ALK+ ALCL cell line. Moreover, this research demonstrates the extent to which binding sites for these AP-1 transcription factors overlap, and identifies many putative novel transcriptional targets for c-Jun/JunB in ALK+ ALCL. Future work will determine whether these are *bona fide* transcriptional targets of these transcription factors, and what role they play in ALK+ ALCL.

## Methods

### Cell lines and culture

The ALK+ ALCL cell line, Karpas 299 (provided by University of Cambridge Enterprise; Cambridge, UK), and the SUP-M2 cell line were obtained from the Leibniz Institute DSMZ-German Collection of Microorganisms and Cell Cultures (Leibniz, Germany). The SR (also termed SR-786) and DG75 cell lines were obtained from the American Type Culture Collection (ATCC). The UCONN-L2 ALK+ ALCL cell line was obtained from Dr. Raymond Lai (University of Alberta). ALK+ ALCL cell lines were grown in RPMI 1640 media (Millipore Sigma) supplemented with 10% Fetal Bovine Serum (Millipore Sigma), 1mM sodium pyruvate, 2mM sodium glutamine, and 50µM 2-mercaptoethanol. In some cases, antibiotics/anti-mycotics (Gibco Thermo Fischer) were added to the media. HEK 293 Tet-On cells stably expressing a doxycycline-inducible NPM-ALK (HEK 293 Tet-On NPM-ALK cells) were a gift from Dr. Raymond Lai and have been previously described (47). These cells were grown in DMEM media (Millipore Sigma) supplemented with 10% Tet system approved FBS (Takara Bio USA), 1mM sodium pyruvate, 2mM sodium glutamine, and 50µM 2-mercaptoethanol. All cells were incubated at 37°C in 5% CO_2_.

### Antibodies and other reagents

The JunB rabbit monoclonal antibody (mAb) (C37F9) used for ChIP-Seq was purchased from Cell Signaling Technology, whereas the c-Jun antibody (H-79) used for ChIP-Seq was from Santa Cruz Biotechnology. The anti-JunB mAb (C11) used in western blotting and EMSA experiments were purchased from Santa Cruz Biotechnology. The anti-Bid mAb (B-3) for EMSAs and anti-β-actin mAb (DM1A) used in western blotting experiments were also purchased from Santa Cruz Biotechnology. Anti-FAM129B antibodies were purchased from Cell Signalling Technology (#5122) and Thermo Fisher (PA5116934). The anti-phospho-FAM129B Ab (S679/683; PA595683) was also purchased from Thermo Fisher. The anti-Erk (137F5), anti-phospho-Erk (T202/Y204; 20G11), and anti-phospho-ALK (Y1278/1282/1283; #3983) were purchased from Cell Signalling Technology. The anti-ALK mAb was from Dako, and the ALK inhibitor, Crizotinib, was purchased from Biovision. The MEK inhibitor, U0126, and doxycycline used for the induction of NPM-ALK expression in the HEK 293 Tet-On cells, were purchased from Millipore Sigma.

### ChIP-Seq experiments

Karpas 299 cells were grown to exponential phase and cell pellets were sent to Active Motif (Carlsbad, CA) for ChIP-Seq analysis. Sequences were aligned to the hg19 build of the human genome and peak calling was done with MACS 2.1.0 (56). Binding sites between c-Jun and JunB were considered to overlap if any if there was any overlap between the two (or more) peaks. Motif enrichment shown in **Fig 1C** was performed with the findMotifsGenome program (v4.8) of the HOMER package (http:homer.ucsd.edu) (24), and represents the top enriched motif when the top 1,000 peaks (+/-200 bp) were examined. The ChIP-Seq data has been deposited in the Gene Expression Omnibus (GEO) database (GSE151413). For analysis of c-Jun/JunB binding sites, BigWig files were uploaded to the Broad Institute’s Integrative Genome Viewer (IGV) (57) (version 2.10.0) and line graphs of mean peak size were examined.

### g:Profiler analysis

Genes associated with c-Jun, JunB, or both c-Jun and JunB were searched using the g:Profiler (38, 39) website (https://biit.cs.ut.ee/gprofiler/gost; version e104_eg51_p15_3922dba) and enriched pathways (adjusted *P* ≤ 0.05 after Benjamini-Hochberg false discovery rate correction) in GO Biological Process, KEGG, and Reactome pathways identified.

### EMSAs

Nuclear extracts from the indicated cell lines were prepared using the Ne-Per nuclear and cytoplasmic kit from Thermo Fischer Scientific. The 5’ biotinylated probe upstream of the *FAM129B* transcriptional start site of the larger isoform, and corresponding non-biotinylated versions, were purchased from Integrated DNA Technologies. The wt sequence was 5’-CAGGCTC<u>TGAGTCA</u>CCAGCTG-3’ and mutant sequence was 5’-CAGGCTC<u>CAGACAC</u>CCAGCTG-3’. EMSAs were performed using the LightShift Chemiluminescent EMSA kit (Thermo Fischer) essentially using the protocol outlined by the manufacturer.

### Luciferase reporter experiments

Nucleotides +50 to -755bp (relative to the transcriptional start site) of the larger *FAM129B* isoform were cloned into the pGL2basic luciferase promoter plasmid (Promega). The AP-1 mutant was constructed by changing the TRE site, TGAGTCA, to CAGACAC. The *GZMB* luciferase construct, and corresponding mutant, have been previously described (17). 2×10^7^ Karpas 299 or DG75 cells were transfected with 10µg of the indicated pGL2 plasmid and 1µg of the *Renilla* luciferase construct. The latter construct was used to control for transfection efficiency. For assays using DG75 cells, 10μg of either control plasmid or pcDNA3.1A expressing Myc-tagged JunB, were also transfected. 24h post-transfection, luciferase activity was measured as previously described (17) using the Dual-Glo Luciferase assay system (Promega) using a Promega GloMax Explorer multimode plate reader.

### shRNA knock-down of FAM129B

The generation of shRNA-containing lentiviral particles and the infection of ALK+ ALCL cell lines has been previously described (18). The shRNA constructs were obtained from Sigma Millipore (control (SHC002); FAM129B shRNA A2 (TRCN0000140763); FAM129B shRNA A3 (TRCN0000122833); JunB shRNA #1 (TRCN0000014943); JunB2-1 shRNA (TCRN0000232083); JunB2-2 shRNA (TCRN0000232084)). Cells were placed into puromycin (Sigma Millipore) selection (0.5µg/ml) 2 days after infection and cells were examined within 2 weeks post-infection.

### Cell lysis and western blotting

Cells were lysed in 1% NP-40 lysis buffer (58) and protein levels were quantified using the Bicinchoninic acid protein assay (Thermo Scientific). Samples were run on SDS-PAGE gels before being transferred to nitrocellulose membranes (Bio-Rad). Blots were blocked in Tris-buffered saline (TBS) containing 5% (w/v) skim milk powder before being probed with primary antibody. Blots were then washed and incubated with Horseradish peroxidase-conjugated secondary Abs and bands were visualized using enhanced reagent (Thermo Scientific) and imaging either on film or an ImageQuant LAS 4000 imager (GE Healthcare Life Sciences).

### BrdU/7-AAD labeling

These experiments were performed as previously described (12). Staining was performed using the BD Pharmingen FITC-BrdU flow kit (BD Biosciences), and cells were stained for 30 minutes with 10µM BrdU. Data was collected on BD LSR Fortessa flow cytometer (BD Biosciences) and analyzed using FlowJo software (Ashland, OR.)

### Statistical analysis

For cell cycle analysis, ANOVA were performed using a Tukey *post-hoc* test. For luciferase reporter assays, paired, one-tailed *T* tests were performed. Results with *P* ≤ 0.05 were considered significant.

## List of Abbreviations

ALK+ ALCL: anaplastic lymphoma kinase-positive, anaplastic large cell lymphoma
AILT/AITL: Angioimmunoblastic T cell lymphoma
ANOVA: analysis of variance
AP-1: activator protein 1
BrdU: bromodeoxyuridine
CRE: cAMP response element
ChIP-Seq: chromatin immunoprecipitation-sequencing
EMSA: electrophoretic mobility shift assay
GO: Gene Ontology
GZMB: Granzyme
B; KEGG: Integrated genome viewer
IGV: Kyoto Encyclopedia of Genes and Genomes
PTCL-NOS: Peripheral T cell lymphoma-not otherwise specified
PH: Pleckstrin Homology
PDGFR-β: Platelet-Derived Growth Factor Receptor-β
PD-L1: program death receptor ligand-1
shRNA: short-hairpin
RNA: siRNA: short interfering
RNA: TRE: TPA responsive elements.

## Acknowledgements

The authors would like to thank Drs. Troy Baldwin and Joel Pearson for providing feedback on the manuscript, and the laboratory of Dr. Raymond Lai for the 293P Tet-ON NOM-ALK cells. Flow Cytometry was performed at the University of Alberta, Faculty of Medicine and Dentistry Flow Cytometry Facility which received financial support from the Faculty of Medicine and Dentistry and Canadian Foundation for Innovation (CFI) awards to contributing investigators.

## Supporting information

**S1 Table Genes associated with c-Jun/JunB binding sites previously demonstrated or suggested to be regulated by AP-1 proteins in ALK+ ALCL**. Genes previously demonstrated or suggested to be regulated by c-Jun/JunB in ALK+ ALCL, and found to be associated with c-Jun and/or JunB binding sites in this study. The listed binding sites are all -/+ 10 kb from the gene and the distance of binding sites from the transcriptional start site is indicated.

**S2 Table Representative genes associated with c-Jun/JunB binding sites important in the pathobiology of ALK+ ALCL, but not previously reported to be regulated by AP-1 proteins**. Genes previously demonstrated to be important in the pathobiology of ALK+ ALCL, and found to be associated with c-Jun and/or JunB binding sites in this study. The listed binding sites are all -/+ 10 kb from the gene and the distance from the transcriptional start site is indicated.

**S3 Table Enriched KEGG pathways for genes associated with c-Jun, JunB, or both c-Jun and JunB binding sites**. Genes associated with c-Jun **(A)**, JunB **(B)**, or both c-Jun and JunB **(C)** binding sites were uploaded to g:Profiler on-line website (38, 39), and searched for enriched pathways in the KEGG database (43, 59). Adjusted *P* values were obtained using a Benjamini-Hochberg correction. In **A** and **B**, the top 10 pathways (based on adjusted *P* value) are listed even though not all pathways had an adjusted *P* value ≤ 0.05. In **C**, all pathways with an adjusted *P* value ≤ 0.05 are listed.

**S1 Fig Gating strategy and representative Karpas 299 and SUP-M2 BrdU/7-AAD plots for results summarized in Fig 7. A**, Gating strategy for analysis of cell cycle. **B**, Representative BrdU/7-AAD blots from the indicated cell lines expressing either control or FAM129B shRNAs.

## References

1. Larose H, Burke GAA, Lowe EJ, Turner SD. From bench to bedside: the past, present and future of therapy for systemic paediatric ALCL, ALK. Br J Haematol. 2019;185(6):1043–54. Epub 2019/01/27.

2. Delsol G, Falini, B., Muller-Hermelink, H.K., Campo, E., Jaffe, E.S., Gascoyne, R.D., Stein, H., Kinney, M.C. Anaplastic large cell lymphoma (ALCL), ALK-positive. In: Swerdlow S.H. CE, Harris N.L., Jaffe E.S., Pileri S.A., Stein H., Thiele J., Vardiman, J.W., editor. WHO Classification of Tumours of Haematopoietic and Lymphoid Tissues. 4th ed. Lyon: International Agency for Research on Cancer (IARC); 2008. p. 429.

3. Pearson JD, Lee JK, Bacani JT, Lai R, Ingham RJ. NPM-ALK: The Prototypic Member of a Family of Oncogenic Fusion Tyrosine Kinases. Journal of signal transduction. 2012;2012:123253. Epub 2012/08/02.

4. Pizzi M, Gaudiano M, Todaro M, Inghirami G. Anaplastic lymphoma kinase: activating mechanisms and signaling pathways. Front Biosci (Schol Ed). 2015;7:283–305. Epub 2015/05/12.

5. Staber PB, Vesely P, Haq N, Ott RG, Funato K, Bambach I, et al. The oncoprotein NPM-ALK of anaplastic large-cell lymphoma induces JUNB transcription via ERK1/2 and JunB translation via mTOR signaling. Blood. 2007;110(9):3374–83.

6. Watanabe M, Sasaki M, Itoh K, Higashihara M, Umezawa K, Kadin ME, et al. JunB induced by constitutive CD30-extracellular signal-regulated kinase 1/2 mitogen-activated protein kinase signaling activates the CD30 promoter in anaplastic large cell lymphoma and reed-sternberg cells of Hodgkin lymphoma. Cancer Res. 2005;65(17):7628–34.

7. Drakos E, Leventaki V, Schlette EJ, Jones D, Lin P, Medeiros LJ, et al. c-Jun expression and activation are restricted to CD30+ lymphoproliferative disorders. Am J Surg Pathol. 2007;31(3):447–53.

8. Shaulian E, Karin M. AP-1 in cell proliferation and survival. Oncogene. 2001;20(19):2390–400.

9. Shaulian E, Karin M. AP-1 as a regulator of cell life and death. Nat Cell Biol. 2002;4(5):E131–6.

10. Chinenov Y, Kerppola TK. Close encounters of many kinds: Fos-Jun interactions that mediate transcription regulatory specificity. Oncogene. 2001;20(19):2438–52.

11. Atsaves V, Lekakis L, Drakos E, Leventaki V, Ghaderi M, Baltatzis GE, et al. The oncogenic JUNB/CD30 axis contributes to cell cycle deregulation in ALK+ anaplastic large cell lymphoma. Br J Haematol. 2014;167(4):514–23. Epub 2014/08/26.

12. Zhang J, Wu Z, Savin A, Yang M, Hsu YR, Jantuan E, et al. The c-Jun and JunB transcription factors facilitate the transit of classical Hodgkin lymphoma tumour cells through G1. Scientific reports. 2018;8(1):16019. Epub 2018/10/31.

13. Leventaki V, Drakos E, Medeiros LJ, Lim MS, Elenitoba-Johnson KS, Claret FX, et al. NPM-ALK oncogenic kinase promotes cell-cycle progression through activation of JNK/cJun signaling in anaplastic large-cell lymphoma. Blood. 2007;110(5):1621–30.

14. Wu Z, Nicoll M, Ingham RJ. AP-1 family transcription factors: a diverse family of proteins that regulate varied cellular activities in classical hodgkin lymphoma and ALK+ ALCL. Experimental hematology & oncology. 2021;10(1):4. Epub 2021/01/09.

15. Foss HD, Anagnostopoulos I, Araujo I, Assaf C, Demel G, Kummer JA, et al. Anaplastic large-cell lymphomas of T-cell and null-cell phenotype express cytotoxic molecules. Blood. 1996;88(10):4005–11.

16. Foss HD, Demel G, Anagnostopoulos I, Araujo I, Hummel M, Stein H. Uniform expression of cytotoxic molecules in anaplastic large cell lymphoma of null/T cell phenotype and in cell lines derived from anaplastic large cell lymphoma. Pathobiology. 1997;65(2):83–90.

17. Pearson JD, Lee JK, Bacani JT, Lai R, Ingham RJ. NPM-ALK and the JunB transcription factor regulate the expression of cytotoxic molecules in ALK-positive, anaplastic large cell lymphoma. Int J Clin Exp Pathol. 2011;4(2):124–33. Epub 2011/02/18.

18. Pearson JD, Zhang J, Wu Z, Thew KD, Rowe KJ, Bacani JT, et al. Expression of granzyme B sensitizes ALK+ ALCL tumour cells to apoptosis-inducing drugs. Molecular cancer. 2014;13(1):199. Epub 2014/08/30.

19. Laimer D, Dolznig H, Kollmann K, Vesely PW, Schlederer M, Merkel O, et al. PDGFR blockade is a rational and effective therapy for NPM-ALK-driven lymphomas. Nature medicine. 2012;18(11):1699–704. Epub 2012/10/16.

20. Atsaves V, Zhang R, Ruder D, Pan Y, Leventaki V, Rassidakis GZ, et al. Constitutive control of AKT1 gene expression by JUNB/CJUN in ALK+ anaplastic large-cell lymphoma: a novel crosstalk mechanism. Leukemia. 2015;29(11):2162–72. Epub 2015/05/20.

21. Pearson JD, Mohammed Z, Bacani JT, Lai R, Ingham RJ. The heat shock protein-90 co-chaperone, Cyclophilin 40, promotes ALK-positive, anaplastic large cell lymphoma viability and its expression is regulated by the NPM-ALK oncoprotein. BMC cancer. 2012;12:229. Epub 2012/06/12.

22. Perez-Benavente B, Garcia JL, Rodriguez MS, Pineda-Lucena A, Piechaczyk M, Font de Mora J, et al. GSK3-SCF(FBXW7) targets JunB for degradation in G2 to preserve chromatid cohesion before anaphase. Oncogene. 2012;32(17):2189–99. Epub 2012/06/20.

23. Zhou H, Zarubin T, Ji Z, Min Z, Zhu W, Downey JS, et al. Frequency and distribution of AP-1 sites in the human genome. DNA Res. 2005;12(2):139–50. Epub 2005/11/24.

24. Heinz S, Benner C, Spann N, Bertolino E, Lin YC, Laslo P, et al. Simple combinations of lineage-determining transcription factors prime cis-regulatory elements required for macrophage and B cell identities. Mol Cell. 2010;38(4):576–89. Epub 2010/06/02.

25. Deaton AM, Bird A. CpG islands and the regulation of transcription. Genes & development. 2011;25(10):1010–22. Epub 2011/05/18.

26. Rodig SJ, Ouyang J, Juszczynski P, Currie T, Law K, Neuberg DS, et al. AP1-dependent galectin-1 expression delineates classical hodgkin and anaplastic large cell lymphomas from other lymphoid malignancies with shared molecular features. Clin Cancer Res. 2008;14(11):3338–44. Epub 2008/06/04.

27. Zamo A, Chiarle R, Piva R, Howes J, Fan Y, Chilosi M, et al. Anaplastic lymphoma kinase (ALK) activates Stat3 and protects hematopoietic cells from cell death. Oncogene. 2002;21(7):1038–47. Epub 2002/02/19.

28. Amin HM, McDonnell TJ, Ma Y, Lin Q, Fujio Y, Kunisada K, et al. Selective inhibition of STAT3 induces apoptosis and G(1) cell cycle arrest in ALK-positive anaplastic large cell lymphoma. Oncogene. 2004;23(32):5426–34. Epub 2004/06/09.

29. Weilemann A, Grau M, Erdmann T, Merkel O, Sobhiafshar U, Anagnostopoulos I, et al. Essential role of IRF4 and MYC signaling for survival of anaplastic large cell lymphoma. Blood. 2015;125(1):124–32. Epub 2014/11/02.

30. Rassidakis GZ, Jones D, Lai R, Ramalingam P, Sarris AH, McDonnell TJ, et al. BCL-2 family proteins in peripheral T-cell lymphomas: correlation with tumour apoptosis and proliferation. The Journal of pathology. 2003;200(2):240–8. Epub 2003/05/20.

31. Rust R, Harms G, Blokzijl T, Boot M, Diepstra A, Kluiver J, et al. High expression of Mcl-1 in ALK positive and negative anaplastic large cell lymphoma. J Clin Pathol. 2005;58(5):520–4. Epub 2005/04/29.

32. Dien Bard J, Gelebart P, Anand M, Zak Z, Hegazy SA, Amin HM, et al. IL-21 contributes to JAK3/STAT3 activation and promotes cell growth in ALK-positive anaplastic large cell lymphoma. Am J Pathol. 2009;175(2):825–34. Epub 2009/07/18.

33. Overbeck TR, Hupfeld T, Krause D, Waldmann-Beushausen R, Chapuy B, Guldenzoph B, et al. Intracellular ATP-binding cassette transporter A3 is expressed in lung cancer cells and modulates susceptibility to cisplatin and paclitaxel. Oncology. 2013;84(6):362–70. Epub 2013/05/22.

34. Hupfeld T, Chapuy B, Schrader V, Beutler M, Veltkamp C, Koch R, et al. Tyrosinekinase inhibition facilitates cooperation of transcription factor SALL4 and ABC transporter A3 towards intrinsic CML cell drug resistance. Br J Haematol. 2013;161(2):204–13. Epub 2013/02/26.

35. Chapuy B, Panse M, Radunski U, Koch R, Wenzel D, Inagaki N, et al. ABC transporter A3 facilitates lysosomal sequestration of imatinib and modulates susceptibility of chronic myeloid leukemia cell lines to this drug. Haematologica. 2009;94(11):1528–36. Epub 2009/11/03.

36. Aung T, Chapuy B, Vogel D, Wenzel D, Oppermann M, Lahmann M, et al. Exosomal evasion of humoral immunotherapy in aggressive B-cell lymphoma modulated by ATP-binding cassette transporter A3. Proc Natl Acad Sci U S A. 2011;108(37):15336–41. Epub 2011/08/30.

37. Saiki Y, Horii A. Multiple functions of S100A10, an important cancer promoter. Pathology international. 2019;69(11):629–36. Epub 2019/10/16.

38. Raudvere U, Kolberg L, Kuzmin I, Arak T, Adler P, Peterson H, et al. g:Profiler: a web server for functional enrichment analysis and conversions of gene lists (2019 update). Nucleic Acids Res. 2019;47(W1):W191–W8. Epub 2019/05/09.

39. Reimand J, Arak T, Vilo J. g:Profiler--a web server for functional interpretation of gene lists (2011 update). Nucleic Acids Res. 2011;39(Web Server issue):W307–15. Epub 2011/06/08.

40. Ashburner M, Ball CA, Blake JA, Botstein D, Butler H, Cherry JM, et al. Gene ontology: tool for the unification of biology. The Gene Ontology Consortium. Nat Genet. 2000;25(1):25–9. Epub 2000/05/10.

41. The Gene Ontology Resource: 20 years and still GOing strong. Nucleic Acids Res. 2019;47(D1):D330–D8. Epub 2018/11/06.

42. Jassal B, Matthews L, Viteri G, Gong C, Lorente P, Fabregat A, et al. The reactome pathway knowledgebase. Nucleic Acids Res. 2019. Epub 2019/11/07.

43. Kanehisa M, Sato Y, Kawashima M, Furumichi M, Tanabe M. KEGG as a reference resource for gene and protein annotation. Nucleic Acids Res. 2016;44(D1):D457–62. Epub 2015/10/18.

44. Old WM, Shabb JB, Houel S, Wang H, Couts KL, Yen CY, et al. Functional proteomics identifies targets of phosphorylation by B-Raf signaling in melanoma. Mol Cell. 2009;34(1):115–31. Epub 2009/04/14.

45. Ji H, Lee JH, Wang Y, Pang Y, Zhang T, Xia Y, et al. EGFR phosphorylates FAM129B to promote Ras activation. Proc Natl Acad Sci U S A. 2016;113(3):644–9. Epub 2016/01/02.

46. Zhou X, Yang F, Zhang Q, Miao Y, Hu X, Li A, et al. FAM129B promoted tumor invasion and proliferation via facilitating the phosphorylation of FAK signaling and associated with adverse clinical outcome of non-small cell lung cancer patients. OncoTargets and therapy. 2018;11:7493–501. Epub 2018/12/01.

47. Young LC, Bone KM, Wang P, Wu F, Adam BA, Hegazy S, et al. Fusion tyrosine kinase NPM-ALK Deregulates MSH2 and suppresses DNA mismatch repair function novel insights into a potent oncoprotein. Am J Pathol. 2011;179(1):411–21. Epub 2011/06/28.

48. Bejjani F, Evanno E, Zibara K, Piechaczyk M, Jariel-Encontre I. The AP-1 transcriptional complex: Local switch or remote command? Biochimica et biophysica acta Reviews on cancer. 2019;1872(1):11–23. Epub 2019/04/30.

49. Prutsch N, Gurnhofer E, Suske T, Liang HC, Schlederer M, Roos S, et al. Dependency on the TYK2/STAT1/MCL1 axis in anaplastic large cell lymphoma. Leukemia. 2019;33(3):696–709. Epub 2018/08/23.

50. Desjobert C, Renalier MH, Bergalet J, Dejean E, Joseph N, Kruczynski A, et al. MiR-29a down-regulation in ALK-positive anaplastic large cell lymphomas contributes to apoptosis blockade through MCL-1 overexpression. Blood. 2011;117(24):6627–37. Epub 2011/04/08.

51. Schleussner N, Merkel O, Costanza M, Liang HC, Hummel F, Romagnani C, et al. The AP-1-BATF and -BATF3 module is essential for growth, survival and TH17/ILC3 skewing of anaplastic large cell lymphoma. Leukemia. 2018;32(9):1994–2007. Epub 2018/03/29.

52. Agnelli L, Mereu E, Pellegrino E, Limongi T, Kwee I, Bergaggio E, et al. Identification of a 3-gene model as a powerful diagnostic tool for the recognition of ALK-negative anaplastic large-cell lymphoma. Blood. 2012;120(6):1274–81. Epub 2012/06/29.

53. Schmidlin CJ, Tian W, Dodson M, Chapman E, Zhang DD. FAM129B-dependent activation of NRF2 promotes an invasive phenotype in BRAF mutant melanoma cells. Molecular carcinogenesis. 2021;60(5):331–41. Epub 2021/03/09.

54. Zeng G, Lian C, Li W, An H, Han Y, Fang D, et al. Upregulation of FAM129B protects cardiomyocytes from hypoxia/reoxygenation-induced injury by inhibiting apoptosis, oxidative stress, and inflammatory response via enhancing Nrf2/ARE activation. Environmental toxicology. 2022;37(5):1018–31. Epub 2022/01/08.

55. Cheng KC, Lin RJ, Cheng JY, Wang SH, Yu JC, Wu JC, et al. FAM129B, an antioxidative protein, reduces chemosensitivity by competing with Nrf2 for Keap1 binding. EBioMedicine. 2019;45:25–38. Epub 2019/07/03.

56. Zhang Y, Liu T, Meyer CA, Eeckhoute J, Johnson DS, Bernstein BE, et al. Model-based analysis of ChIP-Seq (MACS). Genome Biol. 2008;9(9):R137. Epub 2008/09/19.

57. Robinson JT, Thorvaldsdottir H, Winckler W, Guttman M, Lander ES, Getz G, et al. Integrative genomics viewer. Nat Biotechnol. 2011;29(1):24–6. Epub 2011/01/12.

58. Ingham RJ, Raaijmakers J, Lim CS, Mbamalu G, Gish G, Chen F, et al. The Epstein-Barr Virus Protein, Latent Membrane Protein 2A, Co-opts Tyrosine Kinases Used by the T Cell Receptor. J Biol Chem. 2005;280(40):34133–42.

59. Kanehisa M, Goto S. KEGG: kyoto encyclopedia of genes and genomes. Nucleic Acids Res. 2000;28(1):27–30. Epub 1999/12/11.

